# Multi-omic comparative analysis of members of the *Akkermansia* genus reveals species-specific adaptations to growth in mucin

**DOI:** 10.64898/2025.12.30.696933

**Authors:** Agastya Sharma, Maria E Panzetta, Moira Overly, Hyojik Yang, Robert K Ernst, Raphael H Valdivia

## Abstract

*Akkermansia muciniphila* is a commensal, mucophilic anaerobic bacterium that influences human host physiology. Although additional prominent *Akkermansia* species have been identified in humans, their responses to mucin-rich environments remain poorly understood. We conducted a comparative analysis of four representative human isolates: *A. muciniphila, A. biwaensis, A. massiliensis,* and *A. durhamii,* focusing on proteins involved in mucin degradation, cell-surface components, and species-specific secreted metabolites during growth in mucin. Our results reveal unique adaptations of *A. muciniphila* to exploit mucin-rich environments, including higher expression of key mucin-degrading proteins during growth in mucin compared to other *Akkermansia* species. We also demonstrate that *A. muciniphila* expresses a significantly greater number of secreted PEPCTERM proteins, which contribute to host colonization. The expression of pili-associated proteins varied across species, with non-*muciniphila* species producing more predicted pili, suggesting the ability to colonize additional niches. Lastly, we find that small peptides previously linked to host and microbiome modulation in the GI tract are over-represented in the metabolomes of non-*muciniphila* species. Conversely, *A. muciniphila* produces more hydroxylated fatty acids, indicating potential mechanisms for modulating host health. These findings highlight genetic and regulatory mechanisms that may explain *A. muciniphila*’s dominance in the human gut.

## INTRODUCTION

*Akkermansia muciniphila* is a commensal gut bacterium that influences human health (1–7). Its abundance in the human gastrointestinal (GI) tract is linked to positive health outcomes, including lower rates of diabetes, obesity, Crohn’s disease, and ulcerative colitis (1–3). However, high levels of *A. muciniphila* are also associated with the progression of Parkinson’s Disease (4, 5), eczema in children (6), and the risk of developing neutropenic fevers following hematopoietic stem cell transplantations (7). The impact of this bacterium on health is also modulated by diet, as *A. muciniphila* can increase gut permeability under low-fiber diets (8, 9) despite exerting the opposite effect under conventional diets (9, 10).

*A. muciniphila* modulates human health through the end products of mucin metabolism, cell surface components, lipids, and secreted proteins that interface with the mucosal immune system and modulate metabolism. For instance, *Akkermansia-*derived acetate, propionate, and γ-aminobutyric acid, resulting from the breakdown of mucin, can influence the development of intestinal epithelial cells (11, 12) and neuronal signaling (13). *Akkermansia* membrane phospholipids (14–18) and pili proteins (chiefly the pilin protein Amuc_1100) activate TLRs that are linked to the function of epithelial cell junctions (14, 19). In addition, *Akkermansia* mucin metabolism may alter the composition of the microbiota. For example, in co-culture models, *Akkermansia* species can enhance the growth of other mucolytic microbes by producing cobalamin or releasing key sugars such as fucose from mucin (20–22).

Most studies have focused on the *A. muciniphila* isolate Muc^T^/BAA-835, which was first identified in 2004 (23, 24). Analyzing genomes reconstructed from metagenomic sequences, as well as new strains isolated from humans and animals, revealed the presence of additional *Akkermansia* phylogroups (25, 26), which have since been reclassified as separate species. Members of the genus *Akkermansia* with representative human isolates include *A. massiliensis* (AmII) (23, 27), *A. biwaensis* (AmIV) (28), *A. durhamii* (AmVI), and *A. muciniphila* (AmI) (29). Recently, *A. massiliensis* has been reported to exhibit potential probiotic activities, suggesting that *Akkermansia* species in general may share activities that benefit host health (30). *A. muciniphila* is the most commonly detected *Akkermansia* species in human stool as inferred from its prevalence in publicly available metagenomic datasets upon re-analysis focused on newly identified *Akkermansia* species (29, 31, 32). *A. muciniphila* can be further divided into subspecies *muciniphila* (AmIa), which includes the MucT strain, and *communis* (AmIb) (29). *Akkermansia* species vary in their genome sizes, average nucleotide identity (ANI), and predicted metabolic functions (21, 29, 33). Based on metagenomic analysis and quantitative PCR (qPCR) of human stool samples, the relative abundance of *Akkermansia* species can range from undetectable to over 15% of the total microbiota (29, 34). While multiple *Akkermansia* species can coexist, a single species is usually dominant, and the dominant species can vary within an individual over time (33). This suggests that yet-to-be-identified ecological factors influence the presence and abundance of *Akkermansia* in the GI tract.

A common feature of all *Akkermansia* species is their capacity to grow on mucin as the sole source of carbon and nitrogen (24, 33). In particular, for *A. muciniphila,* this ability is essential for its capacity to colonize the GI tract, especially when competing with other members of the microbiota, as demonstrated by mutational analysis (35). We suggest that *Akkermansia’s* adaptations to growth in mucin are important for their capacity to influence metabolism and immunity in human hosts. In this study, we conducted a multi-omic comparative analysis of *A. muciniphila*, *A. massiliensis*, *A. biwaensis,* and *A. durhamii* grown in mucin to identify inherent differences between species that could affect gut colonization and their relationships with host health. We present evidence of species-specific regulation of factors required for mucin degradation and cell surface modification. Additionally, we propose that *A. muciniphila* is uniquely adapted to the gut environment compared to other *Akkermansia* species.

## MATERIALS AND METHODS

### Growth conditions

*Akkermansia* strains were cultured anaerobically at 37°C in a Coy chamber (5% H₂, 5% CO₂, 90% N₂). Strains were grown in porcine gastric mucin (Mucin from porcine stomach Type III, Sialic acid bound 0.5%-1.5%, partially purified powder, Sigma Aldrich cat. no. M1778-100G) or synthetic medium containing soy peptone and threonine (35), supplemented with 5 mM cysteine-HCl when required. The strains used in this study included *A. muciniphila* BAA-835 (NCBI: CP001071), *A. massiliensis* Akk0580 (NZ_CP072044.1), *A. biwaensis* Akk0490 (CP072049.1), and *A. durhamii* RCC_12PD (NZ_CP143889.1), each cultured from verified laboratory stocks (29, 33, 35).

### RNA extraction

Strains were first grown in 2 mL synthetic medium with 5 mM cysteine-HCl for 2 days, then subcultured 1:10 into 9 mL of synthetic or PGM medium and incubated anaerobically to late log phase. Cells were pelleted (15,000 × g, 5 min), washed with Phosphate-buffered saline (PBS), flash-frozen in liquid nitrogen, and stored at −80°C. Pellets were resuspended in 200 µL RNAprotect (Qiagen cat. no. 76104, and RNA was extracted using the Qiagen RNeasy Mini Kit (cat. no. 74104) with 10 µL β-mercaptoethanol per mL RLT buffer. Cells were lysed using ZR BashingBead Lysis Tubes (Zymo, S6003-50) for 5 min on a Qiagen vortex adapter, and on-column DNase digestion (Qiagen, 79254) was performed for 30 min. Extracted RNA was submitted to the Duke Sequencing and Genomic Technologies Core Facility for quality assessment, cDNA synthesis, and sequencing. Sequencing was performed using a NextSeq 1000, capturing 50bp paired-end reads. The resultant sequencing reads were demultiplexed, and adapter sequences were trimmed. Quality was assessed using FastQC 0.11.7 (36). Reads of incorrect length and poor quality were eliminated. Next, reads were aligned to their respective genomes using STAR version 2.7.11a (37), followed by RSEM version RSEM 1.3.3 (38) to ensure correct mapping of multi-aligned reads. RSEM-generated expected counts were used for differential expression analysis using DESeq2 (39). A univariate analysis was used to determine gene expression differences between synthetic and mucin-grown samples of discrete species. Between 98.5% and 100% of predicted open reading frames in all species were detected transcriptionally. Genes were considered differentially expressed if log_2_ fold changes were ≥ 1 or ≤ -1 and p-values were ≤ 0.05.

### Mass Spectrometry Analysis

Cultures were grown as described above, and bacteria were harvested at late log phase. Pellets were sonicated in lysis buffer (150 mM NaCl, 50 mM Tris pH 8, 1% dodecyl-β-maltoside, 1 mM PMSF, 50 mM EDTA) using a Branson Sonifier 450 (10 min, 30 s on/off cycles, power 5). Lysates were clarified by centrifugation (15,000 × g, 30 min, 4°C) and then stored at −80°C until subjected to mass spectrometry analysis. Up to 20 µg of total protein was spiked with 200–400 fmol/µg bovine casein as an internal standard, reduced for 15 min at 80 °C in 5% SDS, alkylated with 20 mM iodoacetamide for 30 min at room temperature, and acidified to 1.2% phosphoric acid. Samples were mixed with 559 µL S-Trap binding buffer (90% methanol/100 mM TEAB), loaded onto S-Trap cartridges (Protifi), digested with sequencing-grade trypsin (20 ng/µL; Promega) for 1 hr at 47 °C, and sequentially eluted with 50 mM TEAB, 0.2% formic acid, and 50% acetonitrile/0.2% formic acid. Eluates were lyophilized and resuspended in 12 µL 1% TFA/2% acetonitrile containing 12.5 fmol/µL yeast ADH.

Quantitative LC–MS/MS was performed on 1 µg of each sample using a Vanquish Neo UPLC (Thermo) coupled to an Orbitrap Astral mass spectrometer (Thermo). Peptides were trapped on a Symmetry C18 column (20 mm × 180 µm; 5 µL/min) and separated on a 1.5 µm EvoSep 150 µm × 8 cm column at 50 °C using a 30 min gradient from 5–30% acetonitrile/0.1% formic acid (500 nL/min). Data-independent acquisition (DIA) was performed with full MS scans at 240,000 resolution (m/z 380–980; AGC 4e5) and 4 m/z fixed windows. MS/MS scans were acquired with AGC 5e4, 6 ms fill time, and 27% HCD energy. Each run lasted ∼45 min. Fmols of protein were quantified by measuring the response factor of the instrument against the intensity value of yeast ADH1 (25fmol spiked into each sample).

Spectra were processed in Spectronaut (Biognosys) using *Akkermansia* species-specific protein databases (protein fasta files of genomes accessed from NCBI with previously noted accession numbers) with reversed-sequence decoys (1% FDR by q-value). Absolute abundances were quantified from MS^2^ fragment ion chromatograms aligned across runs. Differential expression was defined as log_2_ fold changes being ≥ 1 or ≤ -1 and p-values being ≤ 0.05.

### Metabolomics

*Akkermansia* strains were cultured anaerobically at 37 °C on synthetic ± 2.5% PGM agar with 5 mM cysteine. Single colonies were expanded sequentially in synthetic, PGM, or mixed media to late log phase (OD ≈ 0.8–1.0). Cultures (1.5 mL) were centrifuged (15,000 g, 5 min, 4 °C); supernatants (100 µL) were frozen at –80 °C. Metabolite abundances were normalized to cell pellet protein amount per sample. To measure protein concentration, cell pellets were lysed in 200 µL RIPA buffer + protease inhibitor (Thermo 89900; Millipore 04693132001), incubated 45 min at 4 °C, and centrifuged (14,000 rpm, 20 min). Lysates were stored at –20 °C, and protein concentrations were determined using the BCA assay (Thermo A55860).

Supernatants were analyzed, and data processing was performed by Metware Biotechnology Inc. (Woburn, MA) for untargeted and widely targeted metabolomics via UPLC–MS/MS. After extraction with methanol/acetonitrile (4:1 v/v) containing internal standards, the samples were processed according to Metware’s standard pipeline. Untargeted profiling used UPLC (ExionLC 2.0) coupled to a TripleTOF 6600+ (AB SCIEX), and targeted quantification used UPLC coupled to a QTRAP 6500+. Metabolites were identified using the Metware, Metlin, HMDB, KEGG, AI, and MetDNA databases. Quantification employed multiple reaction monitoring (MRM). Abundances were normalized to protein content.

Missing metabolite values were imputed as one-fifth of the minimum per metabolite. Features with QC CV < 0.3 were retained. Differential abundance was tested using Student’s t-test or Wilcoxon rank-sum (FDR-corrected). All analyses were conducted in R v4.5.0, with plots generated in ggplot2 v3.5.2. Full code, package lists, and data are available at https://gitlab.oit.duke.edu/as1114/multi-omic-analysis-of-akkermansia-strains.

### FLAT and FLAT^n^ analysis

#### FLAT protocol

For FLAT analysis, bacterial cells were collected using 10 μL pipette tips as described previously (40). For samples in solution, 1 μL of bacterial suspension was applied directly. One 1μL of citrate buffer (0.2 M citric acid and 0.1 M trisodium citrate, pH 3.8) was added to each spot on a stainless-steel MALDI plate, which was incubated in a humidified glass chamber for 30 min at 110 °C. Plates were cooled, washed thoroughly, and air-dried.

#### Microflex information

Mass spectra for FLAT were acquired in negative-ion mode within the lipid matrix, norharmane (10 mg/mL in 2:1 chloroform/methanol), using a Bruker Microflex LRF MALDI-TOF MS operated in reflectron mode. The instrument used a 337 nm nitrogen laser with the random walk off laser off. Analyses were performed at 70% global intensity; 1200 laser shots were summed per spectrum. Agilent ESI Tune Mix was used for calibration. Data were collected from m/z 1000–3000. Ion transfer settings were: Ion Source 1, 20 kV; Ion Source 2, 16.70 kV; lens, 8.75 kV; reflector, 20 kV. TOF vacuum was maintained below 5.5E-7 mbar. Data processing (baseline subtraction, smoothing, peak picking) was performed in FlexAnalysis (Version 3.4).

#### FLAT^n^ protocol

FLAT^n^ samples were prepared as stated above. Tandem MS was performed on a Bruker MALDI timsTOF operated in “qTOF” mode (TIMS deactivated) with a frequency tripled SmartBeam3D 10 kHz, 355 nm laser. Ion transfer parameters were: funnel1 RF, 440.0 Vpp; funnel2 RF, 490.0 Vpp; multipole RF, 490.0 Vpp; CID energy, 0.0 eV; deflection delta, -60.0 V. Quadrupole MS used ion energy 4.0 eV and an m/z range of 1000–3000. Collision cell activation used 9.0 eV collision energy and 3900.0 Vpp collision RF. Spectra were acquired at a 104 μm laser diameter with beam scan on, 800 laser shots per spot, and 70% and 80% laser power, respectively. MS data were collected in negative ion mode, and isolation widths and collision energies were 4–6 m/z and 100–110 eV. Norharmane (10 mg/mL in 1:2 MeOH/CHCl₃) was used for lipid detection. Data were visualized in mMass (Version 5.5.0) using S/N 3.0, 5% relative intensity, peak height 50, and baseline/smoothing enabled. Fragment assignments were made using ChemDraw Ultra (Version 23.1) and Bruker Compass DataAnalysis.

## Contributions

R.H.V., A.S., and M.E.P. designed the research. A.S. and M.E.P. performed the experiments, wrote the code, analyzed the data, and prepared the figures. M.O., H.Y., and R.K.E. performed the nFLAT analysis and resolved lipid A structures. A.S. and M.E.P. wrote the manuscript. R.H.V. supervised the project. All authors reviewed and edited the manuscript.

## Data Availability Statement

The proteomic, transcriptomic, and metabolomic raw data files are publicly deposited and can be accessed at the following locations: Raw protein result files: PRIDE partner repository (41), dataset identifier PXD070311; Raw metabolomic data: http://dx.doi.org/10.21228/M8WV77; Raw RNASeq data: https://www.ncbi.nlm.nih.gov/bioproject/1347790.

## RESULTS

### Transcriptional, proteomic, and mtabolomic analyses reveal species-specific responses to mucin

We characterized representative isolates of four major human *Akkermansi*a species: *A. muciniphila* (strain Muc^T^/BAA-835), *A. massiliensis* (strain Akk0580), *A. biwaensis* (strain Akk0490), and *A. durhamii* (strain RCC_12PD). The *A. muciniphila* genome is 15-18% smaller than the genomes of the other *Akkermansia* species (29, 33) (**Figure 1A and B**). Shared and unique genes (**Figure 1B**) were identified based on their predicted amino acid sequence similarity using Anvi’o (42, 43). Across all four species, 63-78% of the total predicted ORFs in each genome are conserved throughout the genus, forming the “core” *Akkermansia* genome, while 13%-17% are species-specific. Differences in functional annotation of shared and unique genes between these *Akkermansia* species have been previously documented (29, 33) (**Supplementary Table 1**). To examine species-specific proteins, transcripts, and metabolites that are over-represented during growth in mucin, we cultured *Akkermansia* in either porcine gastric mucin (PGM) or synthetic (mucin-free) media and harvested bacteria or supernatants at late-logarithmic growth phases for transcriptomic, proteomic, and metabolomic analysis (**Figure 1C**).

**Figure 1:**
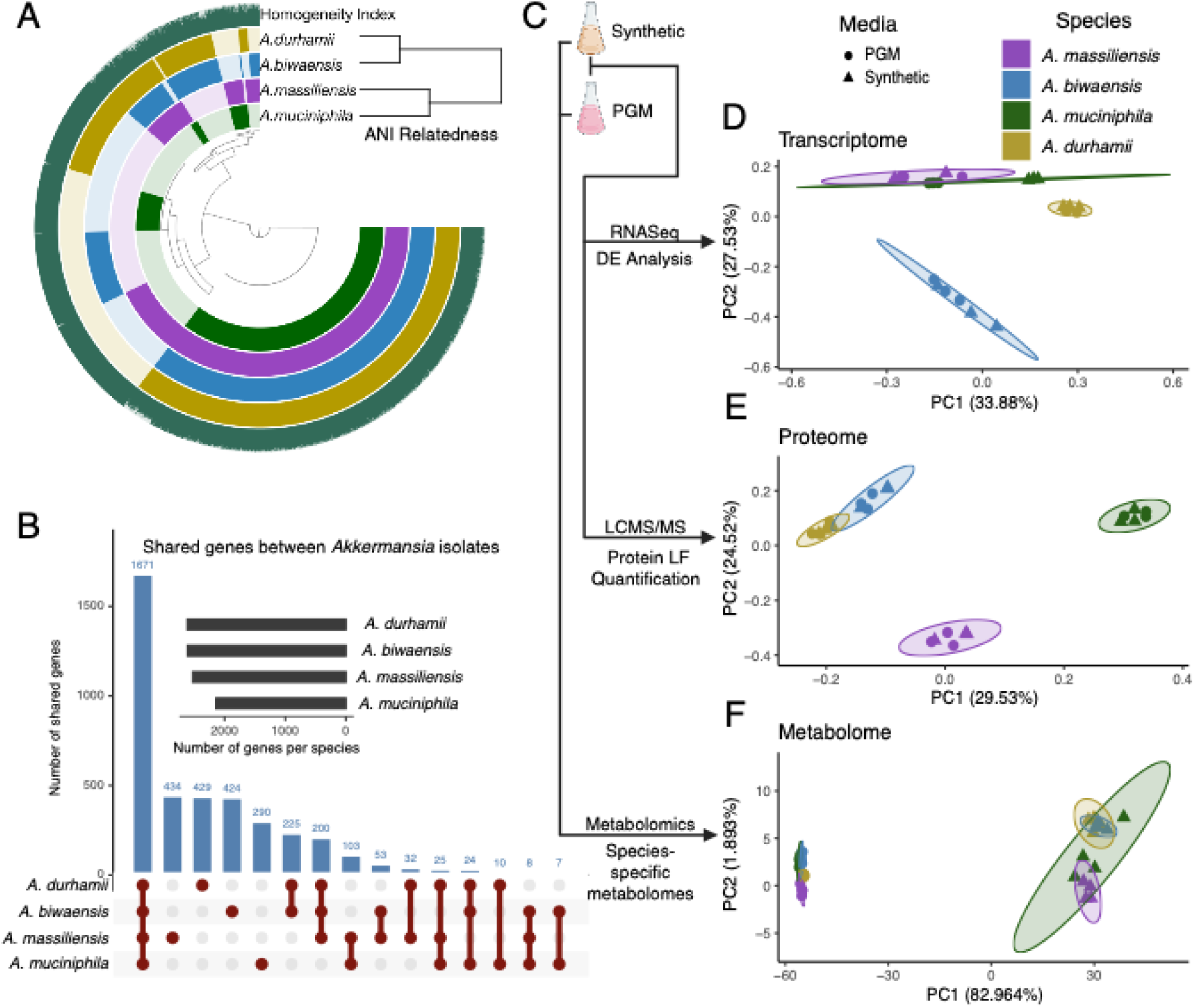
Major *Akkermansia* species display distinct patterns of RNA, protein, and metabolite expression when grown in mucin. **A**. Comparative genomic analysis of representative *Akkermansia* species displays the extent of genetic conservation. The dendrogram shows the phylogenetic relatedness based on average nucleotide identity among the four *Akkermansia* species. **B.** The degree of conservation and uniqueness of genes within this pangenome is highlighted in the UpSet plot. **C.** Schematic of a multi-omic approach to the characterization of the response of *Akkermansia* species to growth in porcine gastric mucin (PGM) or synthetic media. Principal Component analysis of conserved *Akkermansia* RNA transcripts **(D)** and proteins **(E)** expressed, as well as total metabolites identified **(F)** in culture supernatants.

### Transcript

We extracted total RNA from bacterial pellets and processed them for RNA-Seq analysis after removing ribosomal RNA. Reads were mapped to bacterial genes to determine absolute transcript abundance levels, and differential expression analysis identified mucin-responsive genes. Overall, more than 98% of mRNA transcripts matched to predicted open reading frames (ORFs) for each species. A Principal Component Analysis (PCA) based on transcript-per-million values of *Akkermansia* genes conserved among all strains revealed that growth medium, followed by species, were the main factors driving the differences in transcript abundance. The extent of global mRNA changes in response to growth in PGM varied across species (**Figure 1D**), with *A. muciniphila* exhibiting the most significant shift in gene expression.

### Protein

In parallel, we extracted proteins from bacterial pellets, digested them with trypsin, and analyzed all resulting peptides by liquid chromatography-tandem mass spectrometry (LC-MS/MS). We used label-free quantification methods to determine protein abundance during growth in mucin. Between 79% and 85% of proteins that matched predicted ORFs for each species were quantified. To assess whether similar differences were reflected in the proteome, we performed a PCA of conserved *Akkermansia* proteins based on protein abundance (**Figure 1E**). Interestingly, there were no major differences between samples grown in PGM and synthetic media samples for any species, suggesting that the impact of media type on protein abundance is limited. Instead, this analysis revealed that the primary source of variation (PC1) was the species itself, accounting for 29.5% of the variance, despite the analysis being limited to proteins conserved across all species.

### Metabolome

We also performed untargeted metabolomics on the supernatants of *Akkermansia* cultures to evaluate the extent of species-specific metabolites produced during growth in mucin. This analysis detected a total of 1,870 metabolites across all samples. Only 33 metabolites were detected in all species in all media tested (**Supplementary Table 2**); therefore, all metabolites were included in the PCA analysis. Unlike transcriptomic and proteomic data, the metabolic profiles did not distinctly separate one species from another, regardless of the growth medium (**Figure 1F**), indicating that components of the synthetic medium, namely glucose, N-acetylglucosamine, and soy peptone, or mucin, were the primary influences on differences in secreted metabolites. Further analysis of species-specific metabolomes is discussed below.

### Changes in *Akkermansia* protein abundance in response to mucin growth are limited to a narrow range of their proteomes

It was previously reported that approximately 25% of the transcriptome of *A. muciniphila* is differentially expressed in response to PGM (35, 44, 45). In our analysis, both *A. muciniphila* and *A. durhamii* showed significant changes when grown in PGM, with 40.4% and 47.9% of transcripts differentially expressed, respectively. Conversely, the fraction of differentially expressed transcripts was lower for *A. biwaensis* (14.3%) and *A. massiliensis* (7.16%) (**Figure S1A**). Unexpectedly, changes to the proteome of *A. biwaensis* and *A. massiliensis* were more prominent during growth in PGM, with 16.4% and 7.9% of detected proteins affected, respectively, while the opposite trend was observed for *A. muciniphila* (11.2%) and *A. durhamii* (14%).

We found that the overall correlation between transcript and protein differential abundance during growth in PGM versus synthetic media was weak, as shown by Spearman’s rank correlation (**Figure 2A**). To identify subsets of genes with concordant enrichment at both transcript and protein levels, we performed Rank-Rank Hypergeometric Overlap (RRHO) analysis (46) of log2-transformed fold-change values for transcripts and proteins that respond to growth in mucin. RRHO analysis visualizes specific gene subsets with similar expression patterns across larger datasets. This analysis confirmed that highly up-regulated transcripts matched proteins whose levels were also increased during growth in PGM across all species, except for *A. biwaensis.* Conversely, genes with decreased expression and proteins with lower abundance during growth in PGM did not show a correlation **(Figure S2A**). Most genes and proteins with concordant expression belong to the *Akkermansia* core genome, but different subsets were enriched in each species (**Figure S2B**), indicating species-specific regulation of their respective transcriptomes and proteomes during growth in PGM medium.

**Figure 2.**
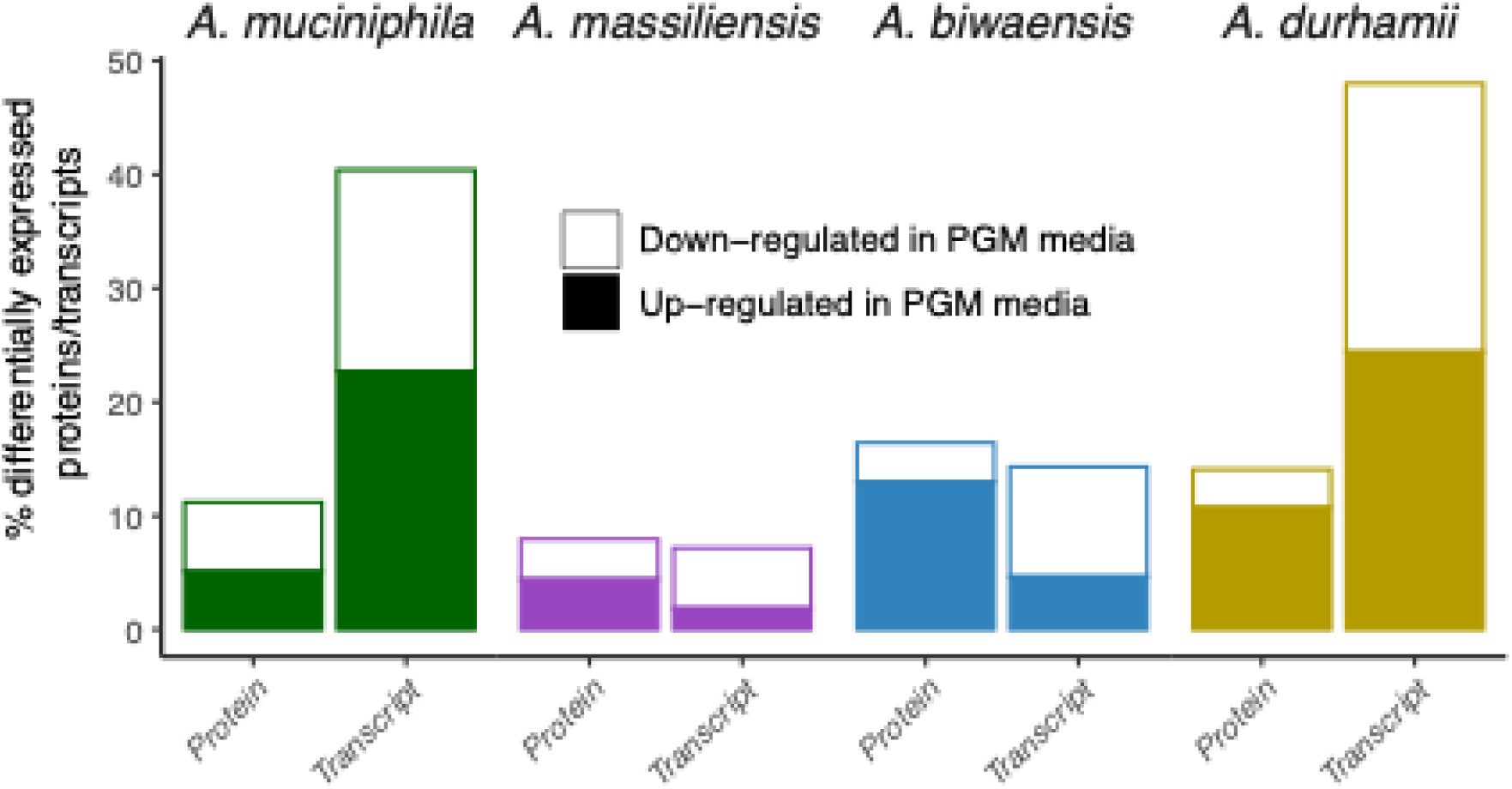
Global protein–transcript correlations are weak across all *Akkermansia* species, and different subsets of conserved proteins are preferentially expressed during mucin growth. **A:** Spearman rank correlation between proteomic and transcriptomic changes (log₂ fold)shows weak or no correlation across all four species. **B:** Rank–rank hypergeometric overlap (RRHO) analysis of pairwise comparisons of proteins expressed during growth in PGM identifies a small set of commonly upregulated core proteins (shown in red) shared across species. Proteins are arranged by fold change from lowest (–) to highest (+) along each axis. The maximal –log₁₀(p) value is shown for each quadrant where overlap is detected. **C:** An UpSet plot summarizing these pairwise RRHO overlaps shows that each species possesses a distinct set of core proteins that increase in abundance during growth in PGM.

To determine whether the same set of conserved *Akkermansia* proteins show an increase in abundance during growth in PGM across different species, we performed a pairwise RRHO analysis of the levels of conserved proteins detected in each species (**Figure 2B**). All *Akkermansia* species showed increased expression of a subset of conserved proteins during growth in PGM, although only 19 were shared across all species (**Figure 2C**). Notably, this was not observed at the transcript level **(Figure S2C)** and therefore, our subsequent analyses will focus exclusively on changes at the proteome level.

### Glycosyl hydrolases are expressed at higher levels in *A. muciniphila* than in other *Akkermansia* species

Multiple glycoside hydrolases (GHs) and endoproteases break down the complex glycan linkages found in mucin (47). GHs are classified into families based on amino acid sequence similarity (48). *Akkermansia* GHs have been identified that target galactose, GlcNAc, GalNAc, sialic acid, and fucose linkages within mucin (47, 49, 50). Unlike other mucolytic microbes, *Akkermansia* processes mucins at the bacterial surface and in the periplasm (35, 47, 51). Mucin-derived monosaccharides and amino acids are then transported into the cytoplasm to fuel *Akkermansia*’s metabolism (35, 47).

We curated a list of predicted GHs and other carbohydrate-active enzymes based on CAZy database annotations (48) among the *Akkermansia* species we tested. Of the 97 GHs identified in the genomes of the *Akkermansia* species, 52 are shared across all genomes and were detected at the protein level in at least one species (**Figure 3A**). All non-*muciniphila Akkermansia* expressed an additional conserved set of 10 GHs. A further 31 GH proteins were expressed in different subsets of *Akkermansia* species. Despite the diversity of GHs, most enzymes belonged to shared GH families (**Figure 3B**), with one *A. biwaensis*-specific GH family represented by a single GH, which was not detectable at the protein level.

**Figure 3:**
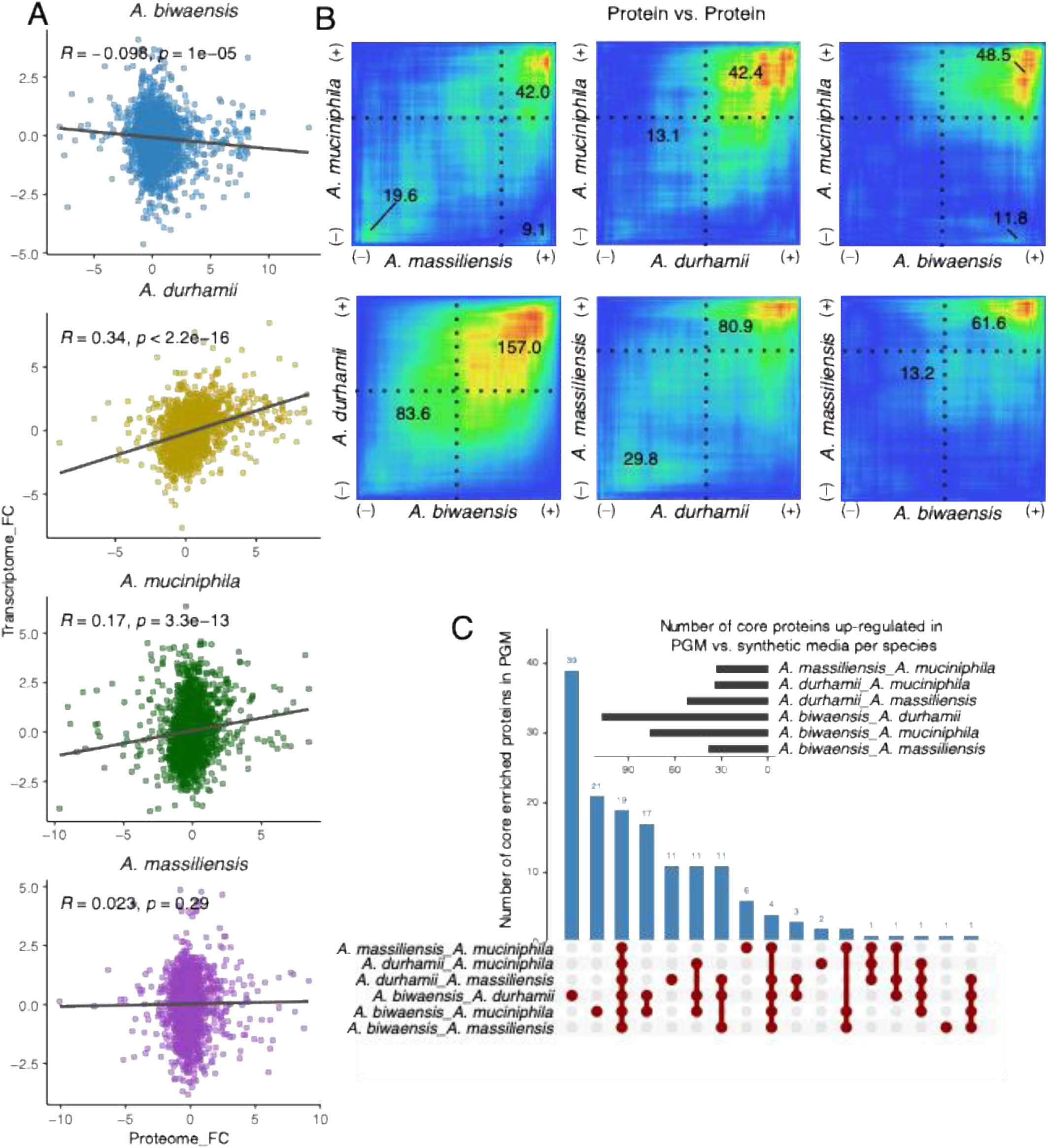
*A. muciniphila* expresses higher levels of glycosyl hydrolases than other *Akkermansia* species. **A-B**: UpSet plot of individual predicted glycosyl hydrolase (GH) enzymes detected by mass spectrometry highlights conserved and species-specific GHs across *Akkermansia* species (**A**). Despite this increased repertoire of enzymes in non-*A. muciniphila* species (>30), GH families are largely conserved across all species (**B**). **C**: PCA of the expression of predicted mucin-degrading proteins indicates that the levels of expression of GH13, GH16, GH18, GH27, and M60 peptidase families distinguish *A. muciniphila* from other *Akkermansia* species. **D:** Total GH abundance (fmol) shows that *A. muciniphila* alone exhibits increased GH abundance during growth in PGM relative to synthetic media. P-values indicate significance of difference in total GH mass between synthetic and PGM media.

We next examined differences in the expression of conserved GHs and mucin-degrading proteases among *Akkermansia* species (**Figure 3C**). A PCA analysis indicated that the main source of variation in mucin-degrading enzyme expression is the growth media, which separated along PC1 (38.19% of total variance). *A. biwaensis, A. durhamii,* and to a lesser extent *A. massiliensis*, display the most significant differences in GH expression when grown in PGM versus synthetic media. In contrast, *A. muciniphila* synthetic and PGM samples cluster more closely together. An analysis of the total abundance of mucin-degrading proteins across all species (fmols of protein) shows that, collectively, only *A. muciniphila* expressed significantly higher levels of GHs in PGM media compared to synthetic media (**Figure 3D**). The next source of variation stems from species-specific trends in GH expression, with non-*muciniphila* species separated from *A. muciniphila* along PC2 (15.42% of the total variance). A biplot analysis to identify which mucin-degrading enzymes drive differences in PC1 and PC2 showed that all GH and protease families aside from GH3 (β-GlcNase) and GH77 (amylase) positively correlated with PC1 (**Figure 3C, inset, Figure S3B**). GHs associated with PC2 (species) highlighted preferences for specific enzyme families with similar functions. Although GHs were not collectively increased in abundance during growth in PGM in non-*muciniphila* species, certain GH families were **(Figure S3C)**. GH27 (α-galactosidase) and M60 metallopeptidase enzymes were significantly increased in PGM media in non-*muciniphila* species, whereas GH2 (β-galactosidase), GH3, GH13 (α-glucosidase), and GH18 (sialidase) enzymes were significantly more abundant in PGM media only in *A. muciniphila* **(Figure S3C)**. Interestingly, *A. muciniphila* GH3 and GH13 enzymes expressed as recombinant proteins do not cleave PGM (47, 48).

### Proteins associated with pili formation are highly expressed in *Akkermansia* species

Several *Akkermansia* proteins have been linked to host immune regulation (19, 52–55). Notably, Amuc_1100 acts as a TLR2 agonist with strong immunostimulatory effects (55). Amuc_1100 has been described as a pili-like outer membrane protein with structural similarity to PilO/PilN of the type IV assembly system (56). It is part of a gene cluster (*Amuc_1098-1102*) that contains additional proteins associated with pilus biogenesis (51). This cluster defines the *MUL2* locus, which is important for mucin acquisition (35) and is conserved among all *Akkermansia* species. We examined the expression of these proteins to determine if there are species-specific patterns. Proteins annotated as pili-like proteins, and the potential machinery involved in pili biosynthesis, are broadly conserved and highly expressed across all species (**Figure 4A**). We observed a higher abundance of predicted Type IV and Type II pili and pseudopili in non-*muciniphila* species compared to *A. muciniphila* in synthetic media (**Figure 4A**). We also noted these proteins were significantly more abundant in PGM media in *A. muciniphila,* though this trend is reversed in non-*muciniphila* species. Certain pili proteins (Amuc_1523-24, Amuc_1583, Amuc_0394, Amuc_0745) are required for *in vivo* colonization by *A. muciniphila* (35), indicating these proteins may play a particularly important role in animals. Previous research has also identified protein P9 (Amuc_1631), Amuc_1409, an aspartic protease (Amuc_1434), and a threonyl-tRNA synthase (Amuc_1732) as important secreted factors that influence host immunity and barrier integrity (52, 54, 57–59). Except for Amuc_1409, all these factors are conserved among *Akkermansia* species and expressed at similar levels (**Figure 4B, Supplementary Tables 3-6**).

**Figure 4.**
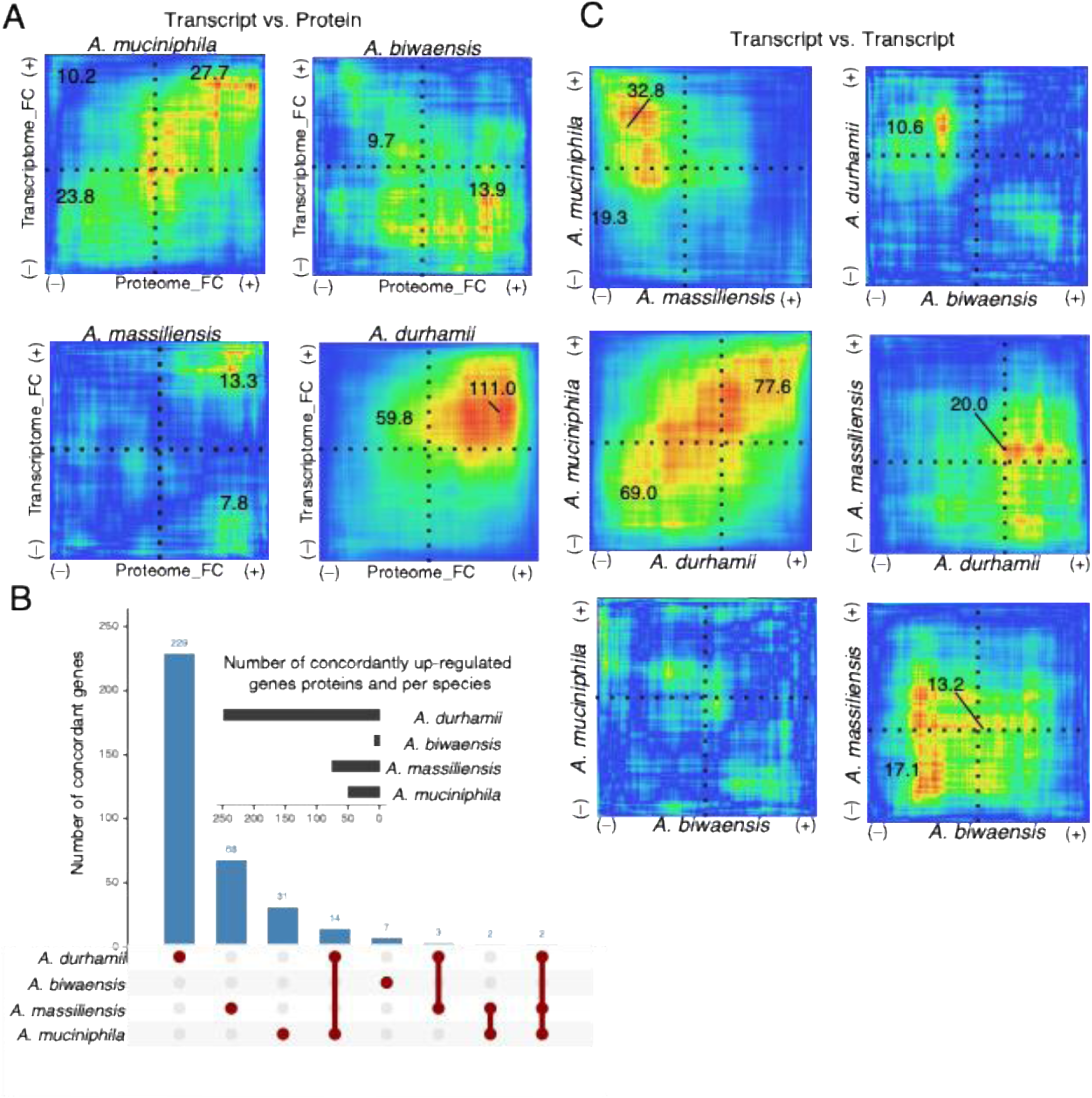
*A. muciniphila* expresses lower levels of pili proteins than other *Akkermansia* species. **A:** Quantified protein abundances (fmol) of proteins annotated as pili assembly components or pili reveal higher expression in *A. muciniphila* compared to non-*A. muciniphila* species. P-values indicate significance of difference in total pili mass between synthetic and PGM media. **B:** Heatmap of absolute protein intensity (log₂-transformed) across species highlights consistent levels of expression of proteins within predicted pili biosynthesis loci, as well as proteins with ascribed immunostimulatory activity in *A. muciniphila*, in all *Akkermansia* strains.

Another key immunomodulatory factor on the cell surface is lipid A, a key component of the outer membrane lipopolysaccharide of Gram-negative bacteria (60). The structure of lipid A varies significantly among bacterial species, and these differences can aid in distinguishing bacteria in clinical samples (61). We determined the lipid A composition of *Akkermansia* species using FLAT^n^ analysis (62). All *Akkermansia* species produce predominantly hexa-acylated lipid A molecules, including two phosphorylated GlcNAc residues, in both synthetic and PGM media (**Figure S4A**). In all *Akkermansia* species, the acyl chains have between 14 and 17 carbon atoms with a branching at the tail, indicating poor recognition by TLR4/MD2 receptors and consistent with reduced pro-inflammatory effects (63). Additionally, mass shifts of *m/z* 123 suggest the addition of phosphoethanolamine (PEtN) groups to Lipid A in all species (**Figure S4A**). We employed FLAT-based tandem mass spectrometry (FLAT^n^) to elucidate the precise molecular architecture of lipid A. This analysis revealed two lipid A molecules at *m/z* 1922.353 and 1824.376 (**Figure S4B and S4C**). Collision-induced dissociation following tandem MS (**Figure S4B and S4C, red lines**) suggests that the loss of a C₂H₄ moiety in the secondary acyl chain at 3′-O is responsible for the mass reduction from *m/z* 1922.353 to 1894.376. Additionally, we identified two additional lipid A ions at *m/z* 2017.330 and 2045.361, which represent the PEtN-modified derivatives of the *m/z* 1894.322 and 1922.353 lipid A species, respectively (**Figure S4D and S4E**). FLAT^n^ confirmed that the PE group is selectively attached to the 1-phosphate position. This modification is known to confer enhanced resistance to antibiotics in Gram-negative bacteria, particularly to colistin (40, 64).

### Amino acid-derived metabolites dominate the metabolomes of *Akkermansia* species

To assess species-specific metabolic profiles derived from identical growth conditions, we conducted untargeted metabolomic analyses of culture supernatants from *Akkermansia* species grown in synthetic medium, synthetic medium supplemented with PGM, or PGM alone. Across all samples, we identified over 1,800 metabolites spanning 24 chemical classes (**Figure 5A; Supplementary Table 2**). *A. durhamii* and *A. biwaensis* shared the greatest number of metabolites across media types, followed by *A. muciniphila* and *A. massiliensis* (**Figure 5E–G**). These shared metabolites largely belong to the same classes, with amino acid-derived metabolites (particularly small peptides <6 amino acids in length, which represent 85% of this category) and organic acid derivatives being the most abundant (**Figure 5A**).

**Figure 5:**
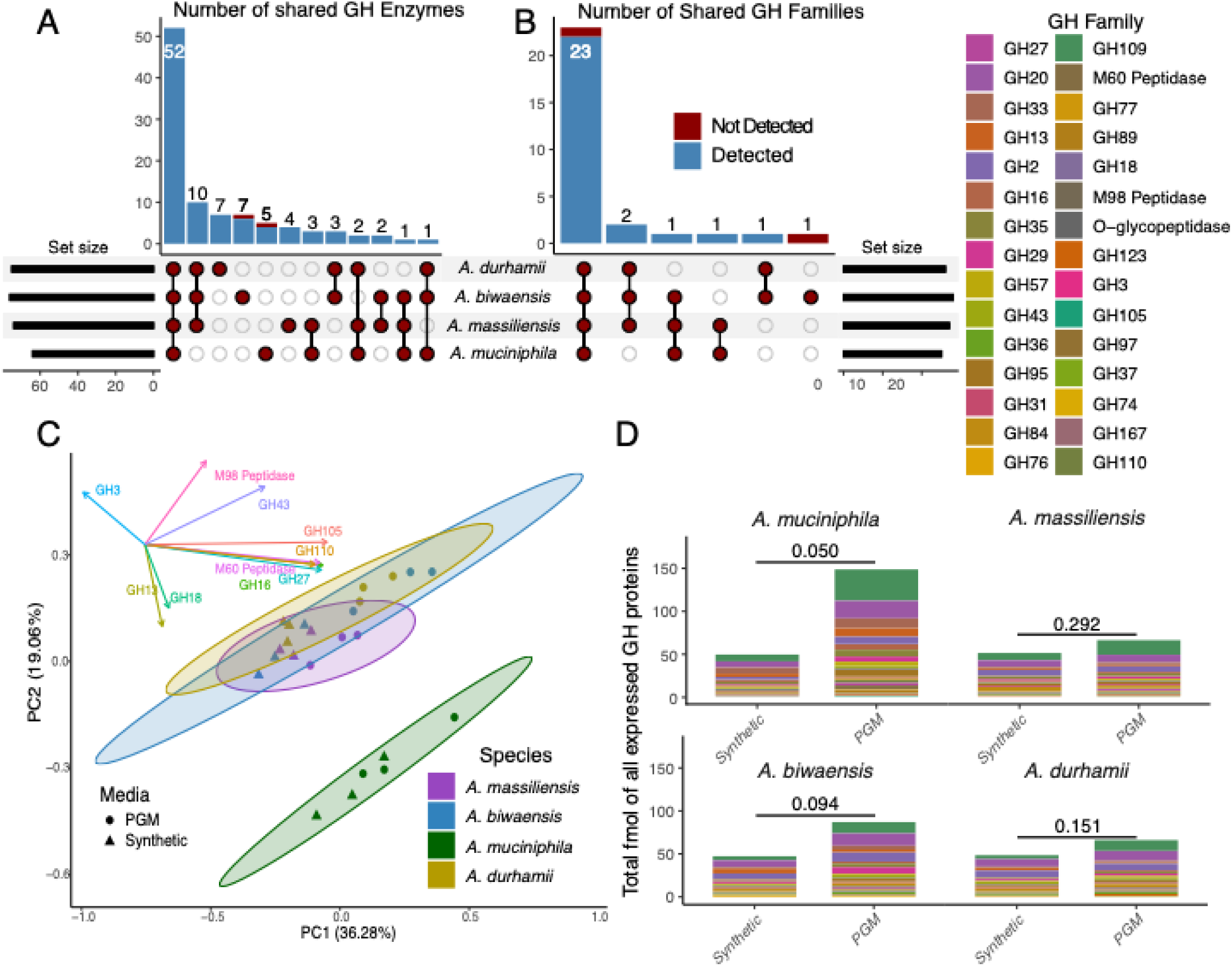
*Akkermansia* species produce unique metabolic signatures when grown in mucin. **A:** Amino acids and their derivatives are the most prominent metabolites produced by *Akkermansia* species across all media types. The ring plot shows the metabolite categories detected in our study. Each color represents a different metabolite class, and the size of each color block indicates the proportion of that class. Principal component analysis reveals little separation between *Akkermansia* species in synthetic (**B**), synthetic + PGM (**C**), and PGM (**D**) media. **E-G:** Metabolites produced in different media (by each strain individually or shared by a combination of strains) categorized by metabolite class. The color of each metabolite class matches that shown in **A**. Amino acid metabolites are significantly over-represented among metabolites produced by non-*muciniphila* species (Fisher’s Exact Test, odds ratio 2.23, p-value 0.0013, FDR 0,03).

Although metabolite profiles were very similar among all strains grown in synthetic medium (**Figure 5B**), metabolite divergence increased with the addition of PGM (**Figure 5C**) and was greatest when PGM served as the only carbon and nitrogen source (**Figure 5D**). The number of unique metabolites also increased as the strains became more reliant on PGM (**Figure 5E**), indicating species-specific adaptations to mucin-based growth.

We further examined metabolites enriched in PGM medium (relative to media-only controls, *p* ≤ 0.05), generating a comprehensive list of metabolites for each species (**Figures 5E–G, S5A**). Among non-*muciniphila*–specific metabolites, amino acid derivatives and small peptides were statistically overrepresented (Fisher’s Exact Test, odds ratio 2.23, p-value 0.0013, FDR 0.03, **Figure 5G**) (65). These amino acid derivatives were among the molecules with the highest fold changes during growth in PGM medium, but not in synthetic medium supplemented with PGM, indicating that the production of these specific metabolites occurs during growth in PGM, not merely exposure to it. Several of these detected small peptides were dipeptides that have previously been shown to have antimicrobial activity (66) **(Figure S5B).** Interestingly, different *Akkermansia* species produced different antimicrobial dipeptides: Trp-Pro (*A. biwaensis*), Pro-Phe (*A. durhamii*), Leu-Thr (*A. massiliensis*), and Tyr-Pro (*A. muciniphila*) (66) (**Supplementary Table 2**). *A. muciniphila* also uniquely generated numerous organic acid derivatives, including hydroxylated fatty acids.

### A large family of PEPCTERM domain proteins characterizes individual *Akkermansia* species

Our analysis of semi-tryptic peptides by LC-MS/MS confirmed that the majority (between 94% and 98%, depending on the species) of *Akkermansia* proteins with predicted secretion signals were cleaved at their predicted sites (67) **(Supplementary Tables 3-6)**. Several of these were found to belong to a class of proteins known as PEPCTERM proteins, previously identified in the outer membrane of *A. muciniphila* (51, 68). PEPCTERM proteins are named for their highly conserved signature C-terminal motif that contains a proline-glutamine-proline tripeptide, followed by a predicted α-helix, and ending in a string of arginine residues (69). The genes encoding PEPCTERM proteins in other bacteria are often linked to exopolysaccharide biosynthesis loci and genes encoding an EpsH/I exosortase system (69). This exosortase is proposed to cleave the PEPCTERM domain after translocation to the periplasm and eventually to the cell surface for secretion (69).

Individual PEPCTERM proteins were not conserved across *Akkermansia* species (**Figure 6A**). Indeed, outside of the PEPCTERM domain, there was little homology at the amino acid sequence level. Because of the uniqueness of PEPCTERM proteins in *Akkermansia* species, we expanded our analysis to include putative PEPCTERM proteins encoded by a broader range of *Akkermansia* genomes. Unsupervised clustering based on the presence or absence of specific predicted PEPCTERM proteins encoded across 89 genomes representing five *Akkermansia* species showed clear distinctions by species, including the two subspecies of *A. muciniphila*, *muciniphila* and *communis* (29) (**Figure S6**). These findings suggest a central role for these proteins in species- and subspecies-specific adaptations to their ecological niches.

**Figure 6:**
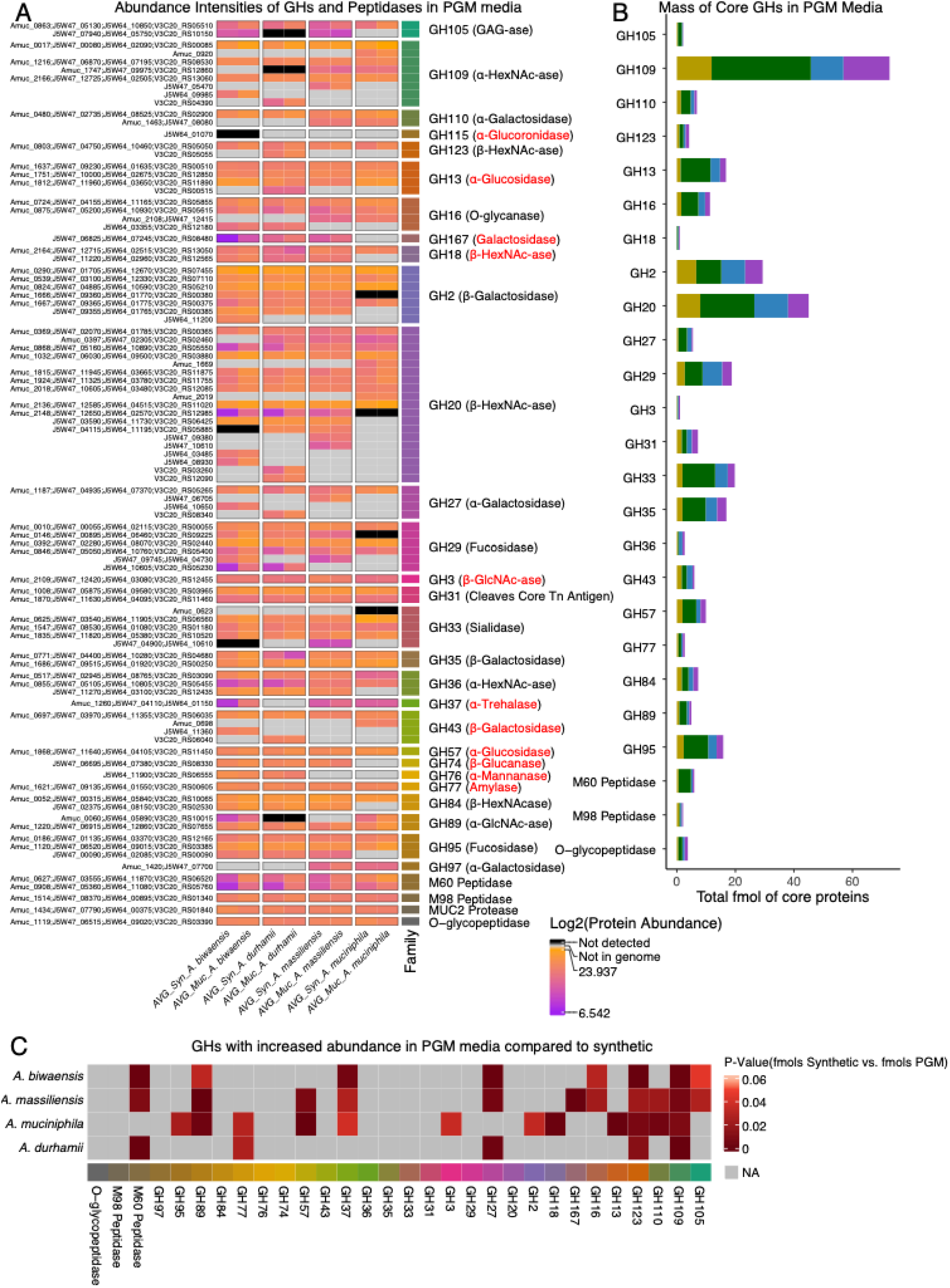
*Akkermansia* PEPCTERM proteins are largely species-specific and are most highly expressed in *A. muciniphila*. **A:** The conservation of PEPCTERM proteins across different *Akkermansia* species is low, as shown by the UpSet plot. Although *A. muciniphila* has the smallest genome, it encodes the most unique PEPCTERM proteins. **B:** PEPCTERM proteins in *A. muciniphila* are expressed at higher levels than in other species (log_₂_-transformed), and the expression of PEPCTERM proteins is significantly increased in non-muciniphila species during growth in PGM media. **C:** Total PEPCTERM abundance (fmol) shows that *A. muciniphila* expresses higher levels of species-specific and pan-*Akkermansia* PEPCTERM proteins during growth in both synthetic and PGM media than other *Akkermansia* species. ** = p < 0.01, *** = p < 0.0001.

We also identified species-specific patterns in PEPCTERM protein expression (**Figure 6B**). *A. muciniphila* produces PEPCTERM proteins at significantly higher levels than other species in both PGM and synthetic media. Additionally, in non-*muciniphila* species, between 18% to 51% of the predicted PEPCTERM proteins were undetectable via mass spectrometry. Six PEPCTERM proteins were conserved across *Akkermansia*, but their overall abundance was much lower than that of species-specific PEPCTERM proteins **(Figure 6C)**. This indicates that the set of predicted PEPCTERM proteins varies greatly in expression and is generally species-specific. For *A. muciniphila, A. durhamii*, and *A. massiliensis*, PEPCTERM protein levels were significantly higher when grown in PGM media, suggesting a role in adapting to the GI tract, and possibly other environments. Conversely, in *A. biwaensis* and PEPCTERM protein abundance did not differ significantly between synthetic and PGM media **(Figure 6B).**

## DISCUSSION

While multiple *Akkermansia* species can colonize the human gut, only *A. muciniphila* has been studied extensively (23). Our results highlight key features that differentiate *A. muciniphila* from other *Akkermansia* species, possibly explaining its high prevalence among human microbiomes.

*Akkermansia* species were clearly distinguishable from one another based on the expression patterns of conserved genes at both the transcript and protein levels. We generally observed low correlation between the transcriptome and proteome across all species (**Figure 2 and S2**). This loss of correlation is likely due to our ability to identify and quantify lower-abundance proteins with new mass spectrometers (>80% of the predicted proteome), thereby increasing the number of proteins whose abundance is less likely to match their respective transcripts. It should be noted that this trend of low concordance between protein and transcript is consistent with reports in other prokaryotic and eukaryotic model systems, where high correlation is usually limited to highly expressed transcripts and proteins (70–74). The relative abundance of conserved proteins in *Akkermansia* proteomes varied by species and was closely aligned with phylogenetic relationships, with *A. durhamii* and *A. biwaensis* sharing the most conserved mucin-responsive proteins (**Figure 2B**). Interestingly, this was not the case at the transcript level, indicating that post-transcriptional regulation plays a significant role in determining final protein abundance (**Figure S2**). Overall, our findings suggest that there are no uniform genus-wide changes in individual protein abundance among *Akkermansia* species in response to growth in PGM. Instead, genus-level trends are observed within specific protein categories. For instance, the transcription of genes encoding GHs in *A. muciniphila* is increased in PGM compared to synthetic media (**Supplementary Tables 3-6**) (35, 44), though this trend was not observed in all species. Correspondingly, the abundance of GHs in *Akkermansia* species, except for *A. muciniphila*, was not significantly higher in PGM media (**Figure 3D**).

Non-*muciniphila* species encode a higher number of GHs compared to *A. muciniphila*. However, these enzymes belong to common GH families (48), suggesting that their overall contribution to mucin glycan degradation is likely conserved. We also noted that several key mucin-degrading GHs were expressed at constitutively high levels in both synthetic and mucin media across all *Akkermansia* species (**Figure S3A**). verall, induction of GH expression in PGM media suggests that *A. muciniphila* is more adapted to break down complex glycosidic linkages in mucin, potentially conferring a competitive advantage when colonizing new mucin-rich environments in the GI tract. In addition, enzymes such as the *A. muciniphila* GH13 (α-glucosidase) lack identified substrates in PGM and are highly expressed during growth on human milk oligosaccharides (75), suggesting that they target non-mucin glycans. The ability to utilize these additional glycans is strain-dependent and may confer advantages within other niches (76).

We also observed conservation in lipid A structures across all species. Recent studies have elucidated the structure of lipid A in *A. muciniphila* (18). Our work indicates that the structure and modifications of lipid A are conserved across *Akkermansia* species and that all are likely poor substrates for TLR4/MD2 recognition, likely due to the terminal branched acyl chains, which is consistent with their reduced pro-inflammatory effects (77). We also identified PEtN as a common modification of *Akkermansia* lipid A across growth conditions (**Figure 4**). This modification is predicted to alter the charge of the bacterial cell membrane, counteracting the positive charge of LPS-targeting antimicrobial peptides (78, 79).

The expanded genomes of non-*muciniphila* species encode species-specific proteins that are differentially abundant during growth in PGM; however, most of these proteins are classified as proteins of unknown function. We predict that some of these proteins contribute to the production of species-specific metabolites during growth in PGM, which may, in turn, provide additional mechanisms for interaction with other microbes, the host, or ecological niches (**Figure 5**). For instance, the dipeptide alanylglutamine, detected exclusively in *A. massiliensis* cultivated in PGM, has been demonstrated to enhance gut barrier integrity following oral supplementation (80). Likewise, several additional *Akkermansia*-derived di- and tri-peptides might serve as signaling molecules capable of modulating their hosts through the epithelial cell transporter PEPT1 (81). Interestingly, the RKH tripeptide, previously identified in *A. muciniphila* cultures and reported to mitigate inflammation in sepsis (82), was not detected in our studies. This is likely due to differences in the growth media used to culture *A. muciniphila*. Indeed, our findings indicate that media composition markedly influences the profile of individual metabolites within a class, with significant differences observed between growth in PGM alone and growth in synthetic media supplemented with PGM (**Figure S5**), suggesting that the *Akkermansia* metabolome is predominantly shaped by carbon and nitrogen sources. Such variability is also anticipated during human colonization, wherein variations in fiber, fat, or protein content in the host’s diet can modify bacterial protein expression and the availability of metabolic precursors to gut bacteria (83).

Another notable group of species-specific factors is PEPCTERM proteins. Based on mutational analysis in *A. muciniphila*, PEPCTERM proteins contribute to colonization in conventionally raised mice but are not necessary for growth in mucin (35). We find that *A. muciniphila* expresses the highest levels of PEPCTERM proteins, even though other *Akkermansia* species encode more of these proteins (**Figure 6**). However, for non-muciniphila species, the more limited repertoire of expressed PEPCTERM proteins is highly induced during mucin growth. We postulate that PEPCTERM proteins likely play a role in adapting to or colonizing mucin-rich environments. For *A. muciniphila*, the higher baseline abundance and greater diversity of PEPCTERM proteins likely enable *A. muciniphila* to colonize and persist more efficiently than other *Akkermansia* species within the mucin layers.

Overall, our findings detail different strategies used by *Akkermansia* species to adapt to mucin-rich environments, with relatively few commonalities in the expression of individual conserved proteins. In fact, we do not detect significant changes in the overall proteome composition during growth in mucin, consistent with specialist organisms adapted to mucus as a dominant niche. Instead, specific categories and families of proteins, such as GHs and PEPCTERM proteins, are the most responsive to growth in mucin, suggesting more tailored responses. *A. muciniphila* appears to be uniquely adapted to mucin-rich environments, which may underlie its apparent dominance in human populations.

## Acknowledgements

We thank Erik Soderblom and the Duke Proteomics Core Facility; Jason Arnold for assistance with RNA extraction and comments on the manuscript, and the Duke Sequencing and Genomic Technologies Core Facility; Katherine Mueller for assistance in genome analysis, and members of the R.H.V. laboratory, Tesia Bobrowski, Elizabeth Thompson, Lauren Davey, and Eric Martens, for their critical review of the manuscript; Pranita Lokinendi for review of the graphical abstract. ChatGPT-5 was used for language improvement and coding assistance. This work was supported by awards from the National Institute of Health (AI142376 and HL170046) and the Howard Hughes Medical Institute Emerging Pathogens Institute, awarded to R.H.V., the support of the University of Maryland School of Dentistry IN-SPIRE grant program, and a National Research Service Award (1F31AI179042-01) awarded to A.S. Schematics were generated in Biorender.

## Declaration of Interest

R.H.V. is a founder of Bloom Science. Bloom Science did not provide funding for these experiments.

---

**Figure S1:**
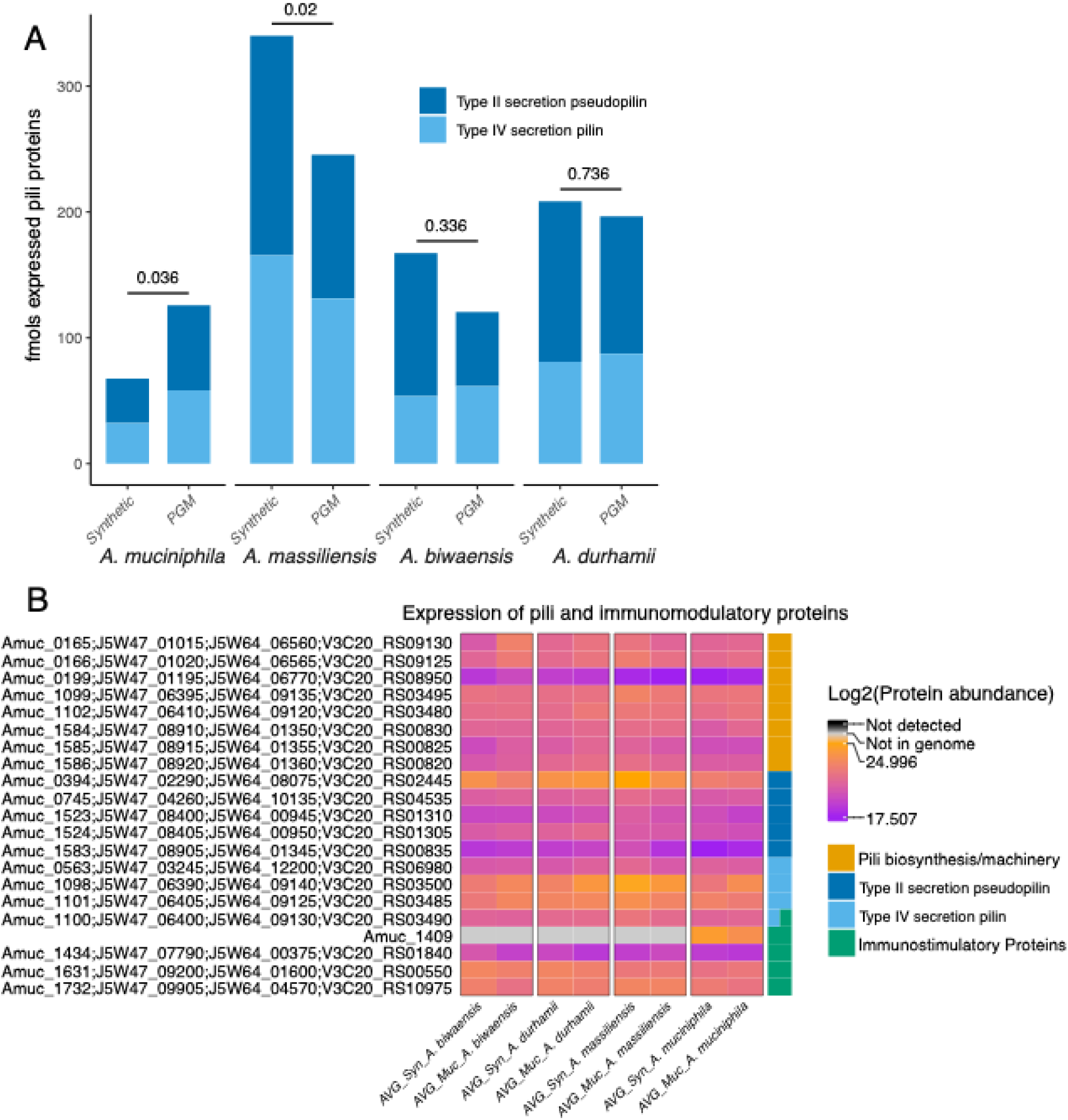
Transcriptional and proteomic changes in *Akkermansia* species in response to different growth media. Proportion of transcripts and proteins whose abundance changes (log_2_ fold change ≥ 1 or ≤ -1 and p-value ≤ 0.05) in response to growth in PGM vs synthetic media, with *A. muciniphila* and *A. durhamii* displaying the most significant transcriptional changes, and *A. biwaensis* and *A. durhamii* displaying the most significant proteomic changes.

**Figure S2.**
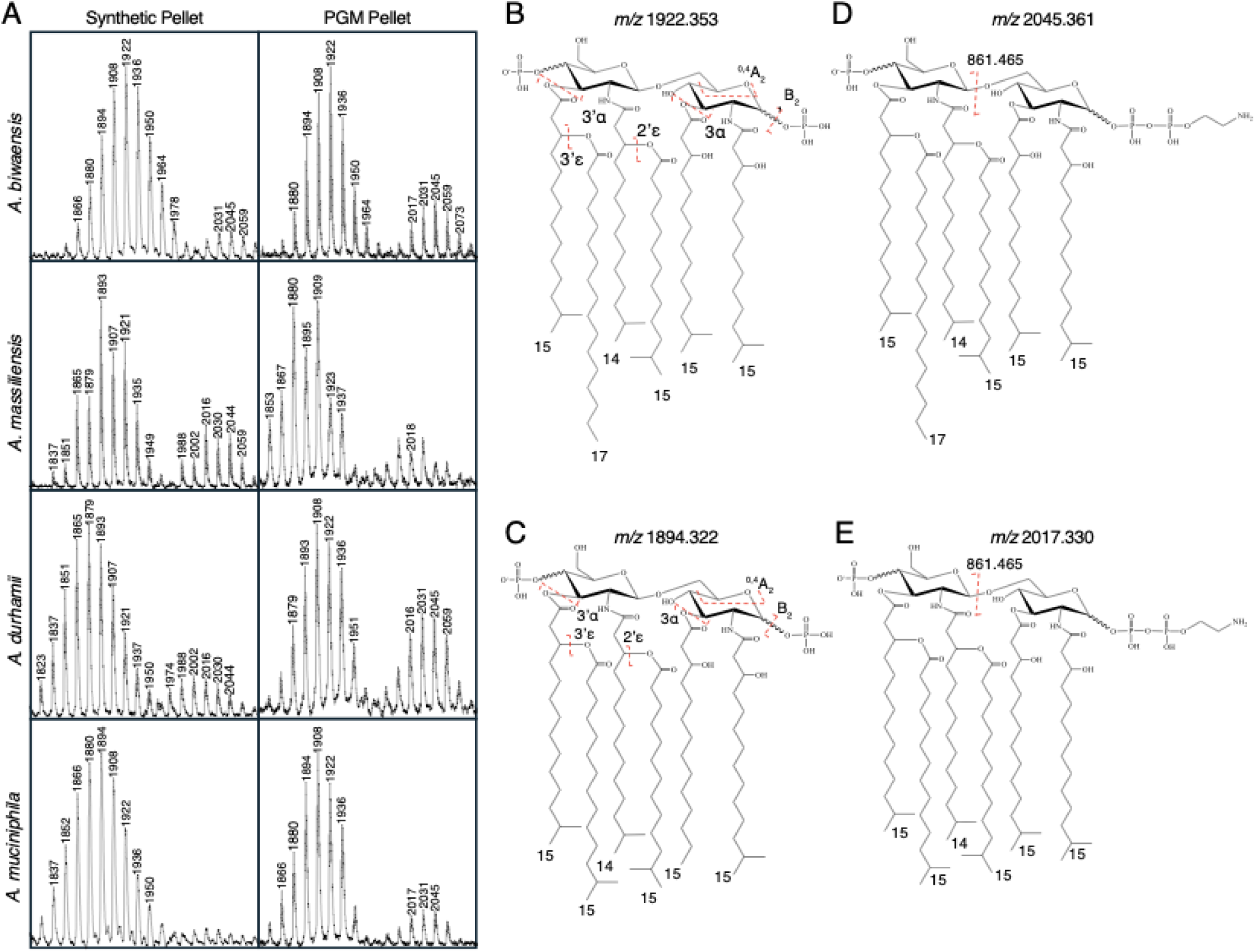
Protein–Transcript concordance within each species contrasts with low transcriptomic concordance across species during growth in PGM media. **A:** RRHO analysis between the transcriptome and proteome of each species shows concordance (shown in red) between highly upregulated transcripts and proteins, except in *A. biwaensis*. Transcripts and proteins are arranged by fold change from lowest (–) to highest (+) along each axis. The maximal –log₁₀(p) value is shown for each quadrant where concordance occurs. **B:** An UpSet plot of genes whose transcript and protein abundance are concordant during growth in PGM. **C:** RRHO analysis between transcriptomes of each species reveals minimal cross-species conservation of gene being expressed (blue), except between *A. durhamii* and *A. muciniphila*.

**Figure S3:**
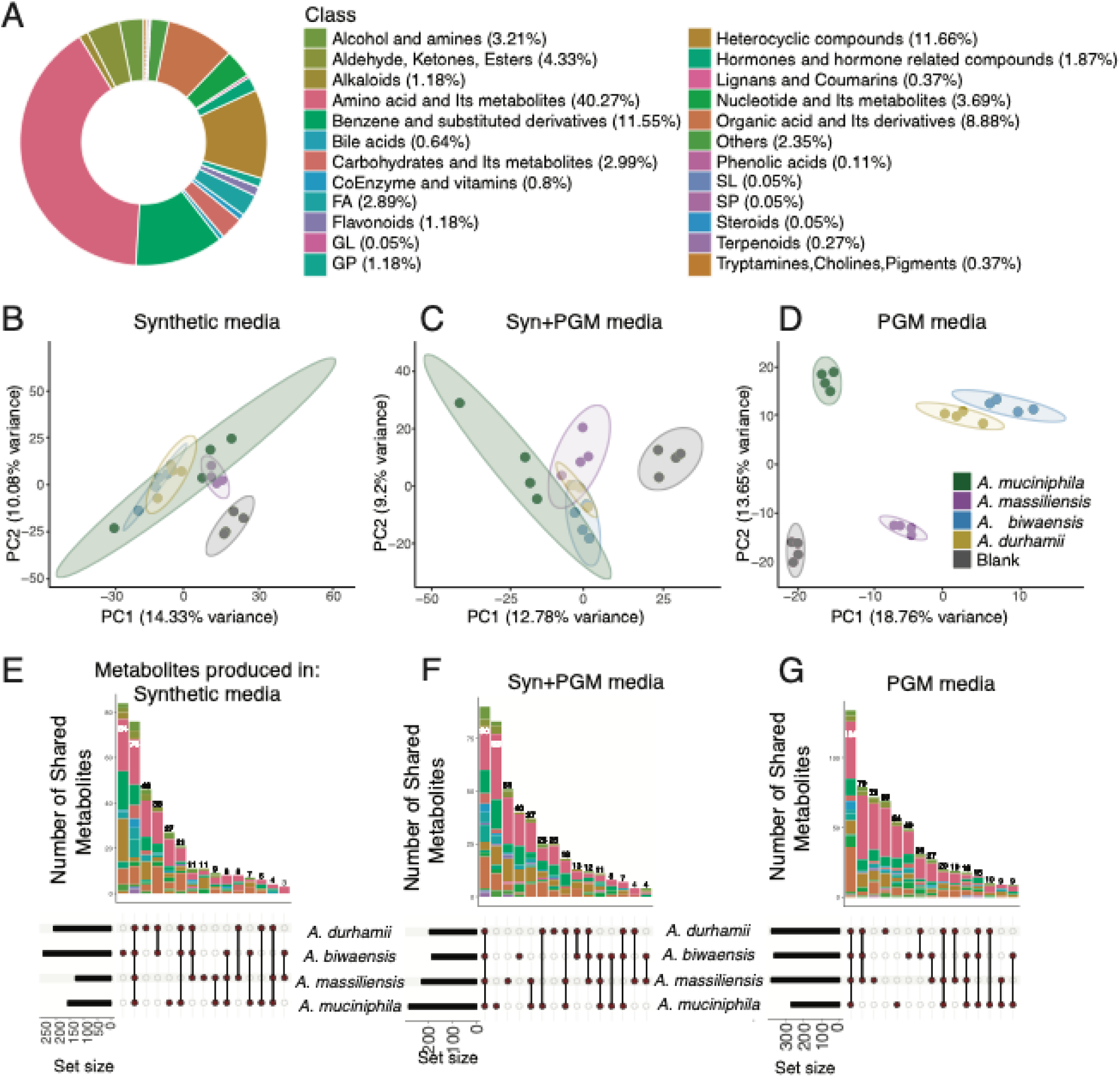
Protein abundance of GHs and GH families in *Akkermansia* species delineates PGM-responsive versus constitutively expressed GHs. **A:** Heatmap of absolute protein (log₂-transformed) for all mucin-degrading proteins quantified by mass spectrometry in the *Akkermansia* pan-proteome reveals GH families that are consistently expressed across species. GH families are annotated by their predicted enzymatic activity; GH families shown in red are not associated with degrading mucin glycans. **B:** α-HexNAc-ase GH109 and β-HexNAc-ase GH20 are the most abundant (fmol) GHs detected during growth in PGM media. **C:** M60 Peptidases and GH27 enzymes are enriched in non-*muciniphila* species, while GH13, GH18, GH3, and GH2 enzymes are enriched in *A. muciniphila.* Heatmap shows p-values of fmols of each GH family in synthetic vs. PGM media.

**Figure S4:**
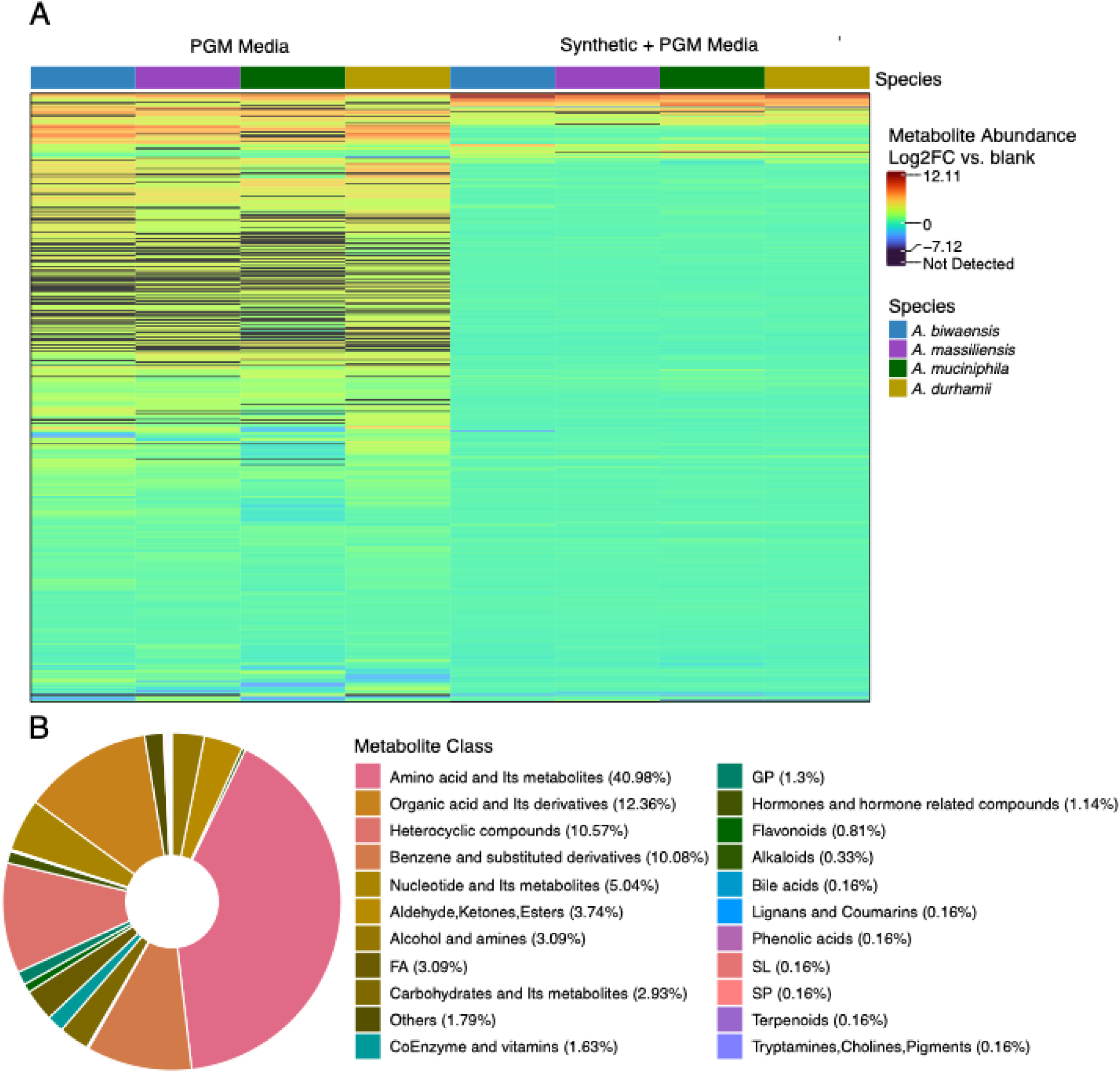
*Akkermansia* species produce hexa-acylated lipid A species that are either bi-phosphorylated or mono-phosphorylated with an added phosphoethanolamine (PEtN) moiety. **A:** Representative GC traces of lipid A purified from cell pellets of all species show the addition of PE in all species and media aside from *A. muciniphila* in synthetic media. **B-E:** Resolved chemical structure of lipid A *m*/*z* 1922.353 **(B)** and *m*/*z* 1894.322 **(C)** molecules from FLAT*^n^*. Resolved chemical structures of PEtN-modified lipid A *m*/*z* 1922.353 and *m*/*z* 1894.322 molecules at *m*/*z* 2045.361 **(D)** and *m*/*z* 2017.330 **(E)**. Red lines indicate bond cleavages induced by collision-induced dissociation (CID) as detected by FLAT^n^.

**Figure S5:**
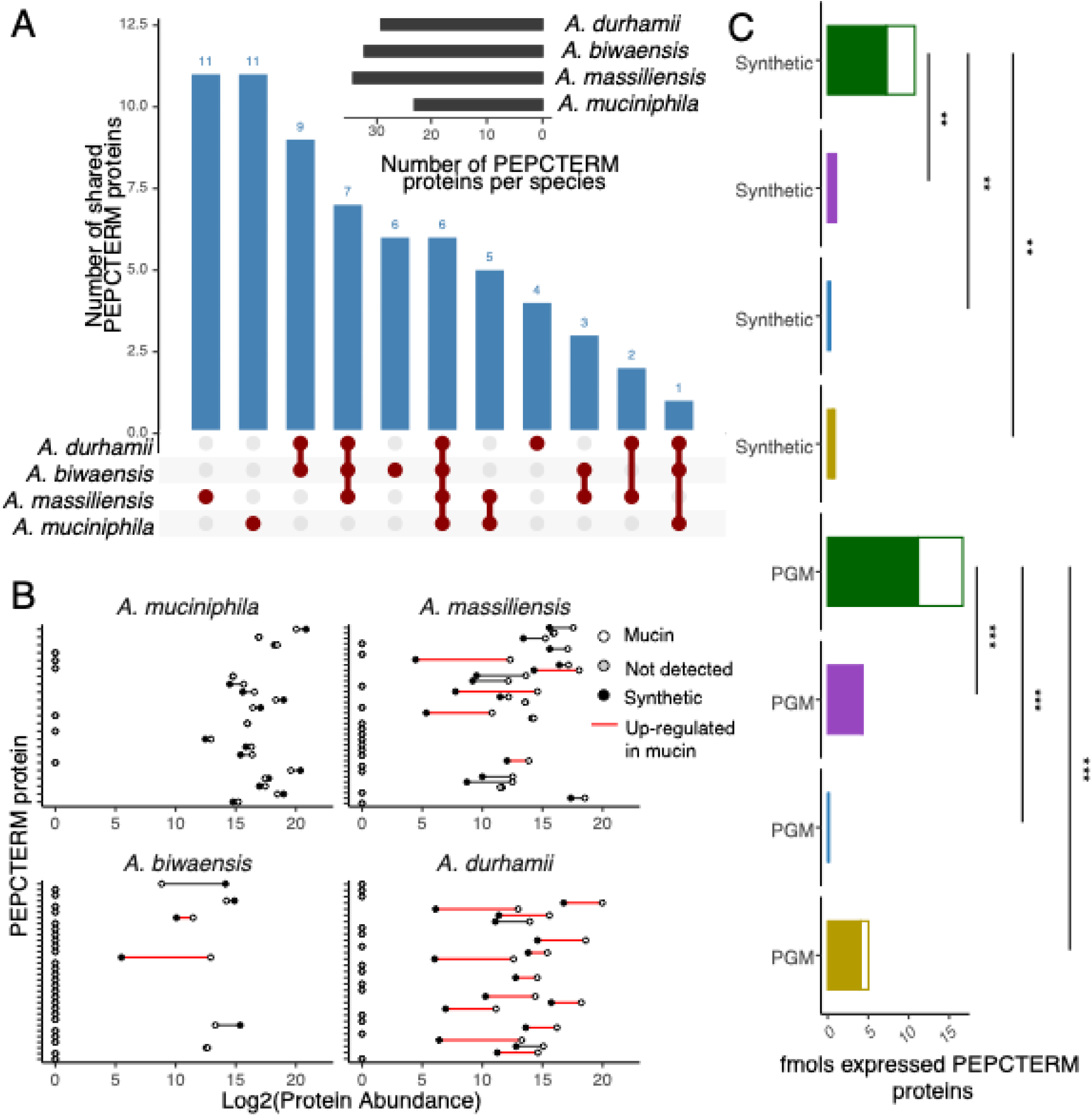
*Akkermansia* produces species-specific metabolites when grown on mucin media. **A:** Metabolites produced in PGM media only are represented in rows, and *Akkermansia* samples in mucin (orange) and synthetic + PGM media (lavender) are represented in columns. Log2-transformed fold changes vs. blank media controls indicate metabolite abundance, and black cells represent metabolites not detected in a given species. **B:** Metabolites identified in (**A)** are separated by class in a ring plot, showing that amino acid-derived metabolites and small peptides, organic acids and derivatives, and heterocyclic compounds are the most abundant class of metabolites produced by *Akkermansia* species during growth in PGM media.

**Figure S6:**
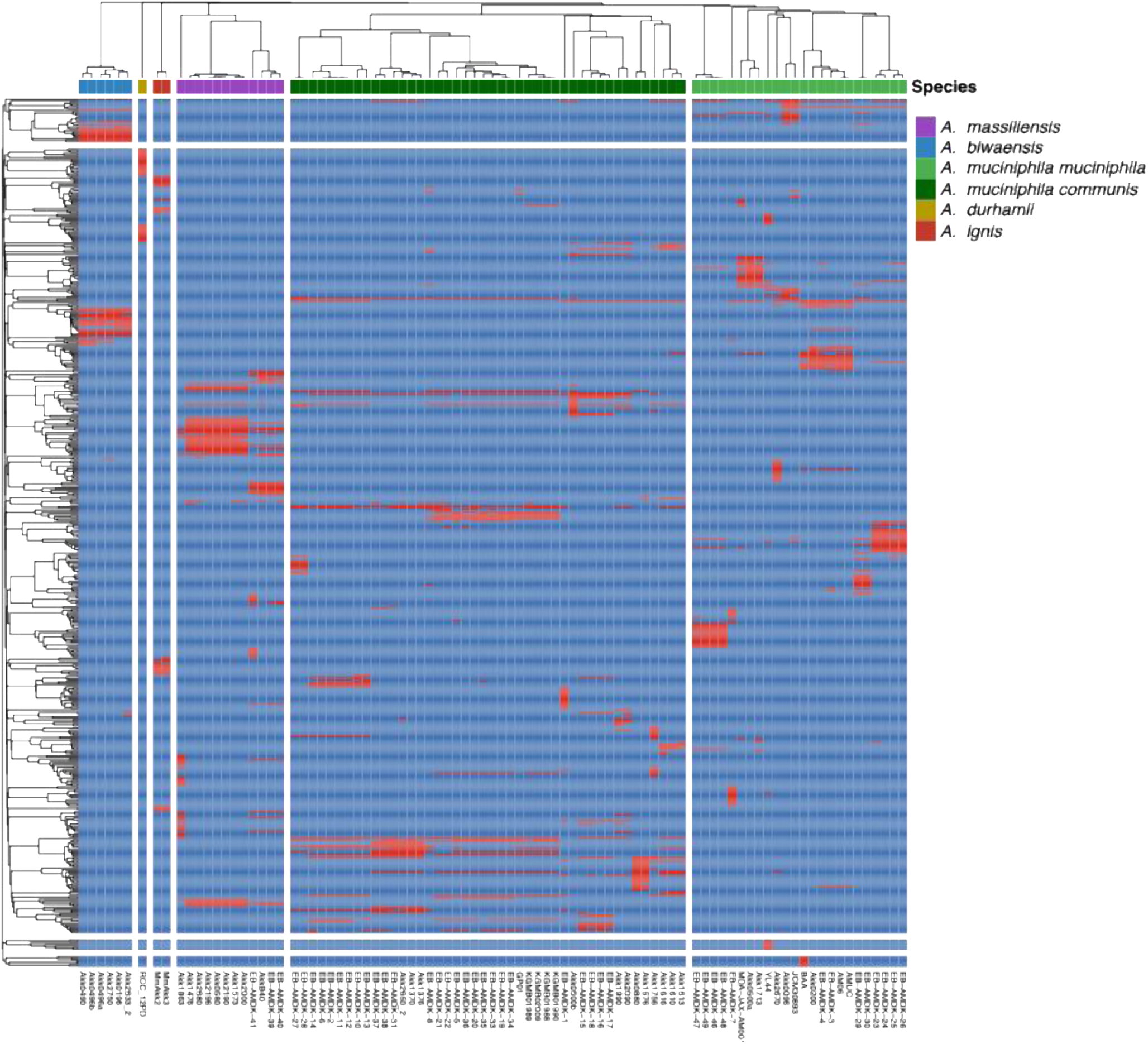
The repertoire of Akkermansia PEPCTERM proteins segregates by species and subspecies. Sequences of predicted 543 PEPCTERM proteins were extracted from 89 *Akkermansia* genomes (labeled columns) and clustered by the presence (red) or absence (blue) of PEPCTERM proteins (rows) identified across all strains.

## Supplementary Table Information

Supplementary Table 1: Gene homology and functional annotations for each predicted gene in *A. biwaensis, A. massiliensis, A. muciniphila,* and *A. durhamii*.

Supplementary Table 2: Raw and imputed metabolites identified in *A. biwaensis, A. massiliensis, A. muciniphila,* and *A. durhamii*.

Supplementary Tables 3-6: Raw proteomic and transcriptomic data for *A. durhamii* (3)*, A. muciniphila* (4)*, A. massiliensis* (5), and *A. biwaensis* (6).

## Notes

http://dx.doi.org/10.21228/M8WV77

https://www.ncbi.nlm.nih.gov/bioproject/1347790

https://www.ebi.ac.uk/pride/

